# Tau-Mediated Cytoskeletal Stabilization Modulates Cell Mechanics and Vulnerability to Mechanical Strain

**DOI:** 10.64898/2026.04.10.717705

**Authors:** Gia Kang, Vineeth Aljapur, Oren E. Petel, Andrew R. Harris

## Abstract

Cells experience mechanical loading across a broad range of loading rates, from low strain rates that are generated during morphogenesis and tissue remodelling, to high and injurious strain rates that are sustained during ventilation-induced lung injury, blast-induced injury, and impact-induced traumatic brain injury. Cell survival under high strain rate loading conditions depends on the ability of the cytoskeleton and plasma membrane to sustain mechanical load without permanent damage. The activity of different cytoskeletal and membrane regulatory proteins could therefore modulate cell susceptibility to injury, but the underlying mechanisms of injury at high strain rate are poorly understood. Tau is a microtubule-associated protein best known for its role in stabilizing microtubules in neurons and as a marker of neurodegenerative disease. Here, we investigated how Tau expression, phosphorylation, and microtubule binding modulates cell viscoelastic behaviour and membrane integrity during high strain-rate uniaxial stretch. We show that Tau expression and de-phosphorylation stabilize microtubules and causes increases in cell stiffness, suppresses cytoskeletal fluidity, and heightens susceptibility to stretch-induced membrane poration. Interestingly, we also find that these effects cannot be explained by microtubule stabilization by Tau alone. Actin architecture acts as a key determinant of injury vulnerability at high strain rate, highlighting the importance of cytoskeletal fluidity and microtubule–actin crosstalk for rapid force dissipation.

**Significance Statement:** Cells must rapidly adapt to mechanical stress during high strain rate deformation. This study shows that Tau, a microtubule-associated protein, modulates cellular mechanics by increasing stiffness, decreasing viscoelastic fluidity, and enhancing susceptibility to membrane poration under rapid stretch. While Tau-mediated microtubule stabilization contributes to these effects, actin architecture and microtubule–actin crosstalk are also critical determinants of injury vulnerability in Tau-expressing cells.

## Introduction

Cells are continually exposed to mechanical strain across a wide range of magnitudes and timescales. While slow deformations accompany processes such as migration, growth, and morphogenesis (strain rate of <0.01 s^-1^), many physiological and pathological contexts impose rapid, high strain rate loading, including tissue stretching (∼10 s^-1^), impulsive fluid shear, compression, and mechanical impact (10 - 40 s^-1^) (1). Rate-dependent cellular injury has been documented in diverse systems. Impact loading of cartilage produces strain rate-dependent chondrocyte death and extracellular matrix damage, contributing to post-traumatic degeneration and osteoarthritis (2–4). Alveolar epithelial cells subjected to rapid stretch or shock-like deformation exhibit increased membrane permeability, oxidative stress, and inflammatory activation, consistent with experimental models of ventilator- and blast-induced lung injury (5–7). Blood cells experience membrane poration and haemolysis under transient high shear, a phenomenon extensively characterized in studies of mechanical circulatory devices and microfluidic constrictions (8, 9). Similarly, endothelial cells exposed to abnormal shear transients display junctional failure, increased permeability, and barrier dysfunction, linking high-rate mechanical forcing to vascular injury and inflammation (10, 11). Neural cells subjected to rapid stretch or shear show comparable rate-dependent injury responses, including membrane poration and cytoskeletal disruption that precede longer-term functional deficits (12–14). These observations indicate that susceptibility to high-rate mechanical injury is widespread but not uniform across cell types, reflecting differences in how effectively cells are adapted to withstand high strain-rate loading. Examining cellular failure under high strain rate deformation therefore offers a means to understand limits of these adaptations and identifying the mechanisms that govern whether cells can accommodate high mechanical loads without injury.

At the cellular scale, the ability to sustain rapid deformation depends primarily on the mechanical properties of the of the plasma membrane, cytoplasm, and cytoskeleton. Force transmission occurs through integrin-cytoskeleton linkages and membrane-cortex attachments (15, 16) that together define cellular viscoelastic properties such as stiffness, fluidity, and strain rate sensitivity (17). When deformation exceeds the system’s capacity for rapid load redistribution, strain concentrates at the membrane-cortex interface, producing transient membrane poration or rupture (18). The cytoskeleton plays a central role in shaping this response. Microtubules provide long-range structural support and resist compression, while the actin cortex forms a contractile, membrane-coupled network that regulates cortical tension. Together, these networks govern how forces are stored, transmitted, or dissipated during rapid loading. Perturbations that alter cytoskeletal dynamics can strongly influence cellular mechanical behaviour. For example, treatment with microtubule-stabilizing agents has been shown to significantly elevate stiffness as measured by atomic force microscopy across multiple cell types (19). In contrast, actin depolymerization or reductions in cortical tension typically soften cells and enhance deformability, while dynamic actin remodelling under strain contributes to strain-softening and energy dissipation (20). These findings suggest that mechanical resilience during rapid deformation depends not on maximal structural stability, but on the ability of cytoskeletal networks to transiently reorganize and redistribute load. However, how distinct cytoskeletal regulatory proteins shape these adaptive responses under high strain rate loading remains poorly understood.

Tau is a microtubule-associated protein that modulates microtubule organization, stability and dynamics in a phosphorylation-dependent manner (21–25). Although Tau is best known for its role in neurons (26) and its relevance to neurodegenerative disease (24, 25), Tau expression has been shown to alter cytoskeletal organization, cell shape, migration, intracellular transport (27, 28), and affect the morphology in non-neuronal cells as well (29). However, how Tau-mediated microtubule stabilization interacts with other cellular structures to influence cellular mechanics during acute, high strain rate deformation remains unclear. Here, we examine how Tau expression, phosphorylation state, microtubule stabilization, and actin architecture collectively shape cellular viscoelastic behaviour and susceptibility to membrane poration under rapid uniaxial stretch. Using atomic force microscopy, live-cell imaging, pharmacological perturbations, and quantitative membrane-permeability assays, we show that Tau expression shifts cells toward a mechanically stiffer, less fluid state that increases vulnerability to rate-dependent membrane failure. Importantly, we find that actin organization is another significant contributor to injury susceptibility in the presence of Tau, revealing that cytoskeletal fluidity, rather than maximal microtubule stabilization, is a key determinant of survival under rapid mechanical loading.

## Results

### Tau expression and phosphorylation increase acute cell injury at high strain rate

We first investigated whether expression of microtubule-associated protein Tau alters cellular sensitivity to rapid mechanical deformation. NIH 3T3 fibroblasts were used as the model system, as they do not endogenously express Tau (30, 31), and have been widely used in cell mechanobiology experiments (32–34). Four human Tau variants were expressed as EGFP fusion constructs: wild-type (WT), a dephospho-mimetic mutant (AP), a phospho-mimetic mutant (E14), and the pathogenic P301L mutation (35). To verify that the transiently transfected Tau is associated with the microtubule cytoskeleton in our system, we imaged 3T3 cells co-transfected with mCherry-tubulin and each Tau construct (Supplementary Fig. 1). Confocal microscopy revealed that all Tau variants exhibited a filamentous distribution that closely followed the endogenous microtubule network, confirming that each Tau construct was association with microtubules.

To assess how Tau expression influences cellular responses to acute mechanical deformation, cells were subjected to single load-and-release cycle of 30% uniaxial stretch at a strain rate of 10 s⁻¹, a loading condition relevant to impact-induced mTBI (36–38). As a metric for cell injury, membrane integrity was quantified by measuring the percentage of propidium iodide-positive (%PI) cells under unstretched and stretched conditions (39). Under unstretched conditions, no significant differences were observed between 3T3 WT cells and Tau-expressing lines or among Tau variants (Fig. 1A). To account for baseline differences in membrane permeability across conditions, we computed the strain-induced change in %PI by normalizing to each treatment’s unstretched baseline (Fig. 1B). 3T3 WT cells differed significantly from all Tau-expressing lines. Among the Tau variants, Tau WT-expressing cells and Tau AP-expressing cells showed the largest increases in stretch-induced PI uptake, and a significant difference was observed between Tau P301L and Tau AP variants. Because these variants are characterized by absent phosphorylation at key sites, this pattern is consistent with a phosphorylation-dependent component to susceptibility to mechanically induced membrane poration.

**Figure 1:**
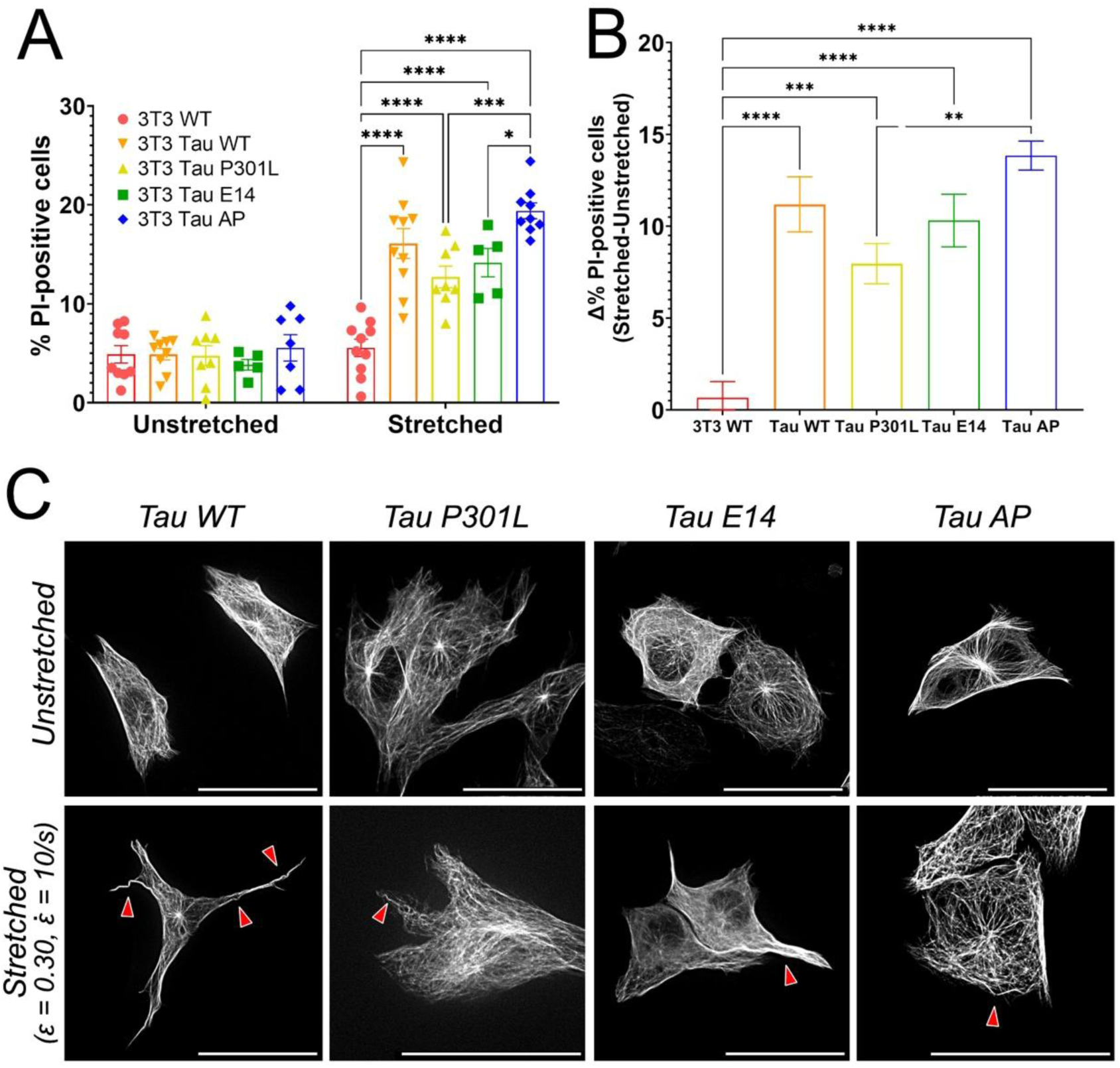
Tau expression increases susceptibility to stretch-induced membrane poration and alters microtubule morphology under load. (A) Percentage of PI-positive cells under unstretched and stretched conditions indicates higher susceptibility to strain across all Tau variants. Membrane poration in the unstretched samples remained low across groups. Following stretch, PI uptake increased across all Tau variants. *Unstretched 3T3 WT: n = 9; Tau WT: p > 0.9999, n = 9; Tau P301L: p > 0.9999, n = 8; Tau E14: p = 0.9718, n = 5; Tau AP: p = 0.9921, n = 7 | Stretched 3T3 WT: n = 10; Tau WT: p < 0.0001, n = 10; Tau P301L: p < 0.0001, n = 8; Tau E14: p < 0.0001, n = 5; Tau AP: p < 0.0001, n = 9; Tau P301L vs. Tau AP: p = 0.0002; Tau E14 vs. Tau AP: p = 0.0226*. Data are presented as mean ± SEM (ns p > 0.05, *p < 0.05, **p < 0.01, ***p < 0.001, ****p < 0.0001) (B) Normalized relative %PI uptake, calculated by subtracting the mean %PI of unstretched controls to reflect strain-induced changes. Strain-induced increases in PI-positive cells differed significantly relative to 3T3 WT. *Tau WT: p < 0.0001; Tau P301L: p = 0.0006; Tau E14: p < 0.0001; Tau AP: p < 0.0001; Tau P301L vs. Tau AP: p = 0.0131.* (C) Representative live-cell images of EGFP-Tau-expressing cells before and after stretch. Post-stretch images reveal localized buckling or waviness of Tau-labelled microtubules (indicated by red arrowheads) without overt network collapse, consistent with reversible structural deformation under load. Scale bar: 50µm.

Live imaging of 3T3 cells expressing EGFP-Tau before and after stretch also showed visible alterations in Tau-bound microtubules (Fig. 1C). Following 30 min of post-stretch incubation, portions of the Tau-labelled network exhibited a wavy or “buckled” appearance (Fig. 1C). However, explicit depolymerization or collapse of the microtubule network was not observed, indicating that the applied strain predominantly induces reversible structural distortion rather than catastrophic filament failure. Our previous work in SH-SY5Y cells (39) similarly indicates that this loading regime reflects transient membrane poration rather than overt cell death.

### Tau expression reduces cellular fluidity and alters viscoelastic creep behaviour

We next sought to find whether differences in Tau phosphorylation state are accompanied by measurable changes in whole-cell mechanical behaviour that could cause the increased stretch-induced membrane poration described above. To test this, we quantified cell stiffness, viscoelastic creep response, and post-load recovery (plasticity) in the untransfected control and Tau-expressing 3T3 cells using atomic force microscopy (AFM) (Fig. 2A-D; see Materials and Methods and (40)). All Tau-expressing conditions exhibited significantly increased cell stiffness relative to untransfected controls (Fig. 2E; p-values presented in figure captions). In contrast, overexpression of mCherry-tubulin did not significantly alter cell stiffness compared to the untransfected control (Supplementary Fig. 2A), indicating that elevated stiffness was not a non-specific consequence of expressing a cytoskeletal fluorescent reporter. Among Tau variants, Tau WT-expressing cells and Tau AP-expressing cells showed the largest increases in stiffness, indicating that Tau expression modulates the mechanical properties of fibroblasts, and that the degree of this effect varies with variant-dependent differences in Tau phosphorylation state.

**Figure 2.**
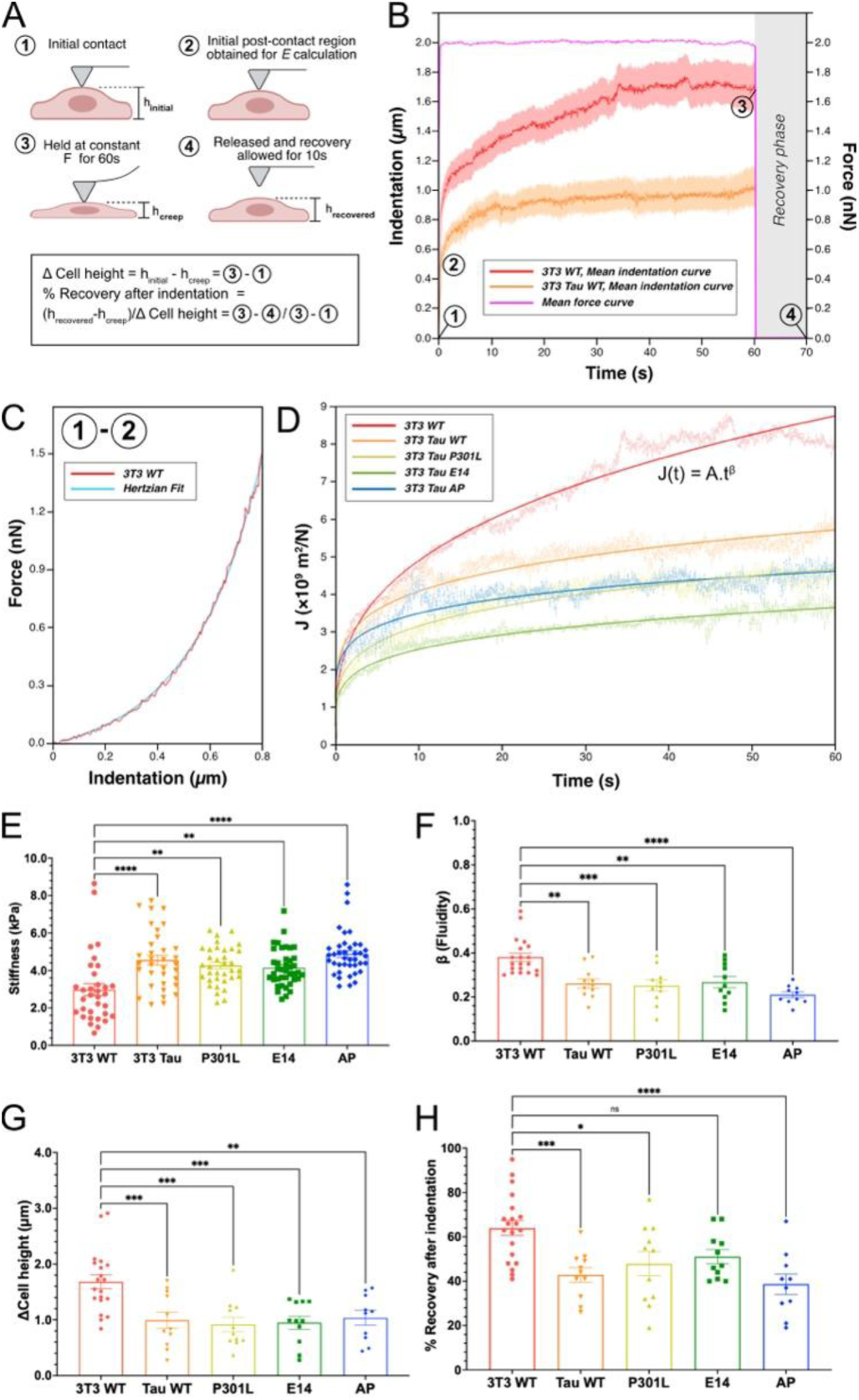
Viscoelastic properties and recovery behaviour of cells expressing Tau variants under mechanical indentation. (A) AFM-based creep assay schematic (2 nN force clamp for 60 s followed by 10 s recovery). (B-D) Representative curves illustrating indentation, constant force hold, and release. Power-law fit of creep compliance J(t)=At^β^ used to calculate fluidity (β). (E) AFM measurements of cellular stiffness reveal significant increases in Tau-expressing cells compared to 3T3 WT cells. *3T3 WT: n = 33; Tau WT: p < 0.0001, n = 34; Tau P301L: p = 0.0010, n = 35; Tau E14: p = 0.0028, n = 40; Tau AP: p < 0.0001, n = 40.* Data are presented as mean ± SEM (ns p > 0.05, *p < 0.05, **p < 0.01, ***p < 0.001, ****p < 0.0001). (F) Fluidity parameter (β) showed reduced cellular fluidity in all Tau-expressing groups compared to WT. *3T3 WT: n = 20; Tau WT: p = 0.0021, n = 11; Tau P301L: p = 0.0007, n = 11; Tau E14: p = 0.0041, n = 11; Tau AP: p < 0.0001, n = 10*. (G) Creep deformation (ΔCell height) showed decreased indentation depth in Tau-expressing cells. *3T3 WT: n = 20; Tau WT: p = 0.0010, n = 11; Tau P301L: p = 0.0002, n = 11; Tau E14: p = 0.0004, n = 11; Tau AP: p = 0.0036, n = 10*. (H) Percentage (%) recovery after force release showed impaired recovery. *3T3 WT: n = 20; Tau WT: p = 0.0004, n = 11; Tau P301L: p = 0.0126, n = 11; Tau E14: p = 0.0807, n = 11; Tau AP: p < 0.0001, n = 10*.

To characterize changes in cellular viscoelastic behaviour with Tau expression, cells were indented with a constant 2 nN force applied for 60 s. From the resulting force-clamp response, we quantified three parameters: (i) the power-law fluidity exponent β (Fig. 2F), (ii) total creep deformation (Fig. 2G), defined as the change in cell height during the 60-s force clamp, and (iii) recovery measured 10 s after load release, expressed as percent height recovery relative to the pre-indentation height (Fig. 2H). Across all Tau variants, β was significantly reduced compared to untransfected controls (Fig. 2F), indicating a shift toward more solid-like behaviour. Consistent with this, Tau-expressing cells exhibited markedly less creep deformation across variants (Fig. 2G). Recovery after unloading was also reduced compared to the untransfected control. Following release of the clamped force, Tau-expressing cells recovered a smaller fraction of their pre-indentation height over the 10-s observation window (Fig. 2H). These results demonstrate that Tau expression increases cell stiffness while reducing cellular fluidity, limiting creep deformation, and impairing short-timescale recovery following load removal which all point to a more solid-like, less dissipative mechanical response during external loading.

### Tau expression and microtubule binding suppresses microtubule network dynamics

To assess whether Tau association with microtubules is correlated with changes in microtubule flexibility and dynamic behaviour, we performed two complementary quantitative assays examining (i) Tau binding dynamics and (ii) microtubule network fluctuations in living cells. First, we quantified Tau binding dynamics using fluorescence recovery after photobleaching (FRAP) (see Materials and Methods), and the initial recovery rate following photobleaching was fitted to estimate Tau turnover on microtubules. Tau E14-expressing cells and Tau P301L-expressing cells exhibited faster fluorescence recovery than Tau WT-expressing cells, indicating more rapid microtubule-binding dynamics. In contrast, Tau AP-expressing cells recovered substantially more slowly, consistent with slower microtubule-binding dynamics. These recovery kinetics align with the established relationship between Tau phosphorylation state and microtubule-binding affinity, in which increased phosphorylation is associated with reduced microtubule-binding affinity (41, 42). To further assess whether Tau directly affects microtubule dynamics in living cells, we quantified temporal fluctuations of the tubulin network using mCherry-tubulin-expressing 3T3 cells with or without co-expression of EGFP-Tau. For each cell, we collected 120-s time-lapse image sequences and computed the coefficient of variation (CoV) of pixel intensities, normalized to the maximum fluorescence signal to account for differences in expression levels. CoV maps were generated on a per-cell basis, and the fraction of pixels exceeding a defined threshold was used as a quantitative measure of microtubule fluctuations (see Materials and Methods; Supplementary Fig. 2B; Supplementary Video). As a positive control, microtubules were pharmacologically stabilized with paclitaxel (43, 44). Paclitaxel treatment markedly reduced tubulin fluctuations relative to control (Fig. 3B). Tau WT-expressing cells showed a similar suppression of microtubule dynamics relative to control, indicating that Tau binding is sufficient to attenuate filament fluctuations.

**Figure 3.**
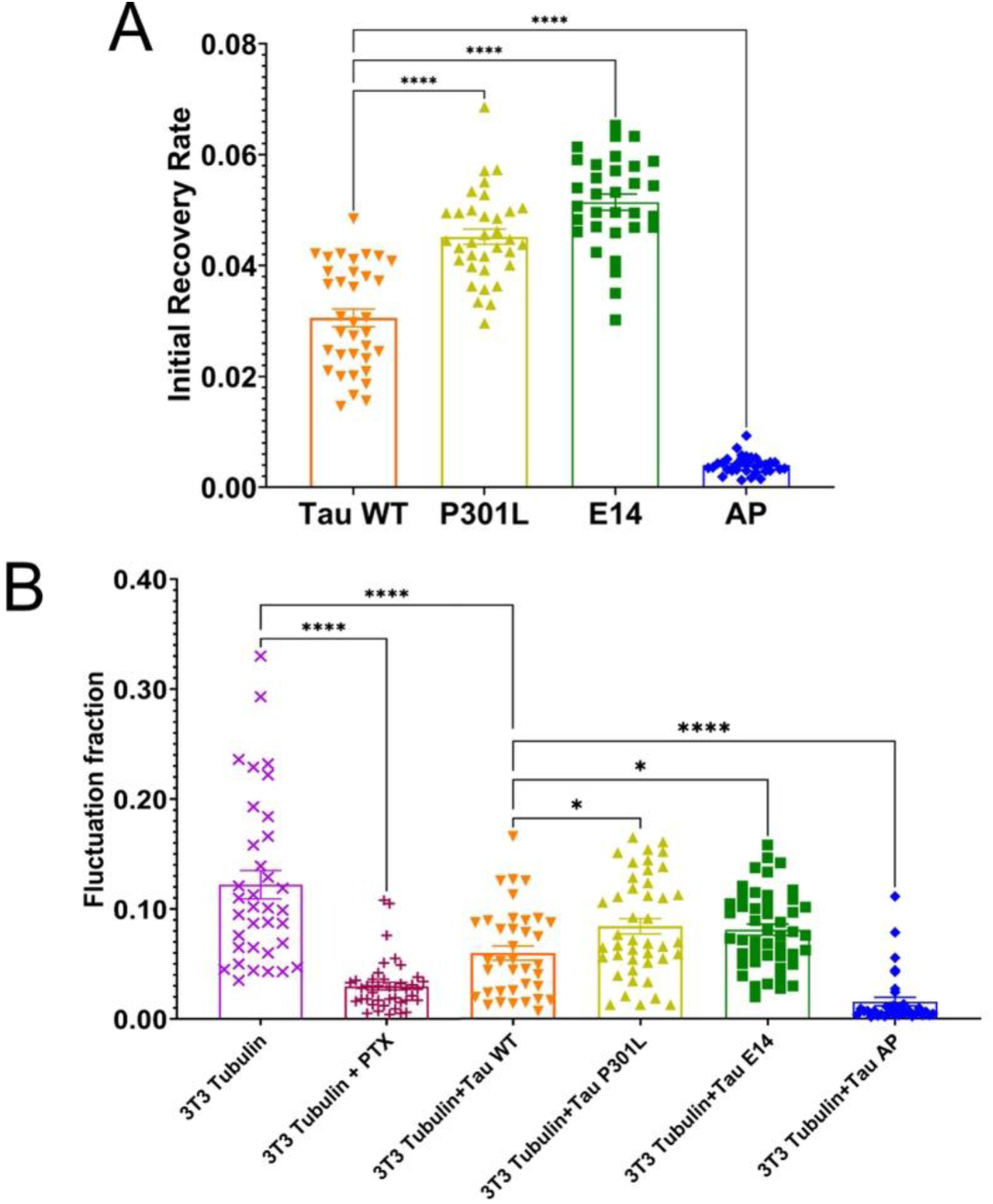
Tau binding suppresses microtubule network dynamics in a phosphorylation-dependent manner. (A) Fluorescence recovery after photobleaching (FRAP) recovery rates for EGFP-tagged Tau variants, illustrating variant-dependent differences in Tau turnover on microtubules. *Tau WT: n = 35; Tau P301L: p < 0.0001, n = 34; Tau E14: p < 0.0001, n = 32; Tau AP: p < 0.0001, n = 34.* Data are presented as mean ± SEM (ns p > 0.05, *p < 0.05, **p < 0.01, ***p < 0.001, ****p < 0.0001) (B) Quantification of microtubule fluctuation fraction derived from time-lapse imaging of mCherry-tubulin. Paclitaxel treatment and Tau WT expression similarly suppressed microtubule fluctuations relative to control. Tau E14- and Tau P301L-expressing cells exhibited higher fluctuation levels than Tau WT-expressing cells, whereas Tau AP-expressing cells showed the strongest suppression. *3T3 Tubulin: n = 37, Paclitaxel: p < 0.0001 vs. 3T3 Tubulin, n = 37; Tau WT: p < 0.0001, n = 37; Tau P301L: p = 0.0104 vs. Tau WT, n = 43; Tau E14: p = 0.0263, n = 47; Tau AP: p < 0.0001, n = 36*.

Consistent with the variant-dependent differences observed in Tau binding kinetics, Tau variants also produced distinct effects on microtubule fluctuation behaviour. Cells co-expressing mCherry-tubulin and Tau E14 or Tau P301L exhibited higher levels of tubulin fluctuation compared to cells expressing Tau WT (Fig. 3B), indicating a more dynamically active microtubule network. In contrast, Tau AP-expressing cells showed marked suppression of microtubule fluctuations relative to Tau WT-expressing cells, yielding the lowest fluctuation levels among all variants tested. These differences mirror the FRAP-derived recovery rates, in which Tau E14 and Tau P301L displayed faster turnover and Tau AP showed slowed recovery and indicate that Tau phosphorylation state modulates microtubule network dynamics in a graded manner rather than producing a binary stabilized or destabilized state.

### GSK-3β inhibition affects membrane poration

Tau phosphorylation state is a common target for neurodegenerative disease treatment (45, 46). Hyperphosphorylation of Tau can lead to neurofibrillary bundle formation, and several drugs targeting GSK-3β (upstream of Tau phosphorylation) have been developed. We therefore hypothesized that pharmacological reduction of Tau phosphorylation could increase Tau binding and increase susceptibility to damage during stretch. To evaluate how Tau phosphorylation state influences susceptibility to mechanical injury, we quantified propidium iodide uptake in Tau WT-expressing 3T3 cells following treatment with the GSK-3β inhibitor tideglusib (TDG). Under unstretched conditions PI positivity remained low (Fig. 4A), indicating minimal baseline toxicity. Surprisingly, TDG treatment reduced stretch-induced membrane poration compared to untreated

**Figure 4.**
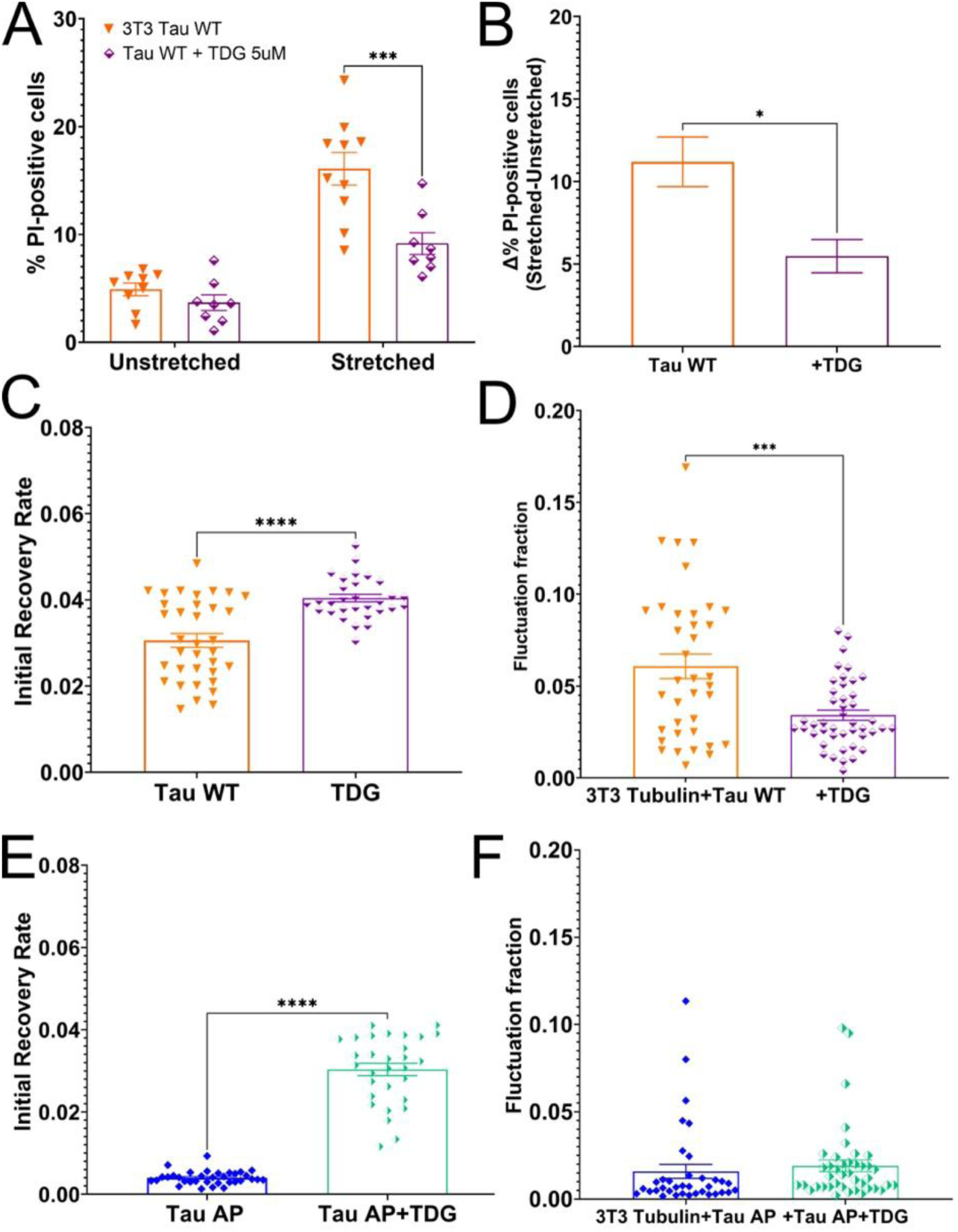
Tau dephosphorylation modulates injury susceptibility and Tau-microtubule binding dynamics. (A) Percentage of PI-positive cells under unstretched and stretched conditions for 3T3 cells expressing Tau WT, with or without treatment with the GSK-3β inhibitor tideglusib (TDG, 5 μM). TDG treatment reduced stretch-induced PI uptake in Tau WT-expressing cells. *Unstretched Tau WT: n = 9; Tau WT + TDG: p = 0.9936, n = 8 | Stretched Tau WT: n = 10; Tau WT + TDG: p = 0.0001, n = 8.* Data are presented as mean ± SEM (ns p > 0.05, *p < 0.05, **p < 0.01, ***p < 0.001, ****p < 0.0001) (B) Normalized relative %PI uptake, calculated by subtracting the mean %PI of unstretched controls to reflect strain-induced changes, highlighting attenuation of stretch-induced membrane poration following TDG treatment. *Tau WT + TDG vs. Tau WT: p = 0.016*. (C) Initial FRAP recovery rate of EGFP-Tau WT in untreated and TDG-treated cells, showing increased recovery following TDG treatment. *Tau WT: n = 35; Tau WT + TDG: p < 0.0001, n = 30*. (D) Microtubule fluctuation fraction measured in 3T3 cells co-expressing tubulin and Tau WT, with and without TDG treatment. *Tau WT: n = 37; Tau WT + TDG: p = 0.0003, n = 44*. (E) Initial FRAP recovery rate of Tau AP before and after TDG treatment, demonstrating increased recovery following TDG treatment. *Tau AP: n = 34; Tau AP + TDG: p < 0.0001, n = 30*. (F) Corresponding microtubule fluctuation fraction in Tau AP-expressing cells with and without TDG; no significant difference was observed following TDG treatment. *Tau AP: n = 36; Tau AP + TDG: p = 0.9995, n = 41*.

Tau WT-expressing cells (Fig. 4B) even though it de-phosphorylates Tau, as confirmed by fluorescence quantification of phospho-Tau in TDG-treated cells (see Supplementary Fig. 3A). To determine whether the reduction in PI uptake could be explained by altered Tau-microtubule binding, we examined Tau dynamics using FRAP. In Tau WT-expressing cells, TDG treatment increased the initial recovery rate compared to untreated controls (Fig. 4C), indicating faster Tau turnover on microtubules. However, this occurred concurrently with a decrease in tubulin fluctuation levels (Fig. 4D). These findings suggest that TDG treatment dissociates Tau turnover kinetics from microtubule fluctuation state in Tau WT-expressing cells, pointing to additional mechanisms linking Tau binding, microtubule dynamics, and susceptibility to stretch. A similar divergence was observed in Tau AP-expressing cells: TDG treatment increased the Tau AP recovery rate (Fig. 4E), whereas microtubule fluctuation levels remained strongly suppressed and were not altered by TDG (Fig. 4F). Because Tau AP cannot undergo phosphorylation-dependent regulation, the observed increase in Tau turnover cannot be attributed to changes in Tau phosphorylation state. Instead, these findings indicate that TDG can alter Tau binding dynamics through a phosphorylation-independent mechanism while leaving the constrained microtubule fluctuation state in Tau AP-expressing cells largely unchanged. These results show that GSK-3β inhibition via TDG reduces stretch-induced membrane poration through a mechanism that cannot be contributed solely to Tau phosphorylation-dependent microtubule binding.

### Actin Organization Feeds back on Tau binding dynamics

This unexpected dissociation motivated further investigation into non-Tau pathways influencing cytoskeletal mechanics and membrane vulnerability under rapid mechanical loading. In the context of tideglusib, GSK-3β functions upstream of small GTPases that control actin polymerization and organization (47–49), which are major contributors to cellular mechanical response (31–33). Therefore, to determine whether actin organization influences Tau-microtubule interactions, we measured Tau binding dynamics while pharmacologically perturbing the actin cytoskeleton. We used a panel of pharmacological agents that target distinct actin regulators and first verified their effects on actin through live-imaging (Fig. 5A) and stress fiber density analysis in LifeAct-GFP-expressing 3T3 cells (Fig. 5B; see Materials and Methods, (53), and Supplementary Fig. 4). NSC23766 was applied to inhibit Rac1 (Rac1-i), selectively perturbing lamellipodial dynamics without directly targeting stress fibers. Y-27632 was used to inhibit ROCK (ROCK-i) and thereby suppress Rho-mediated actomyosin contractility and reduce stress fiber formation. To directly manipulate filament stability, we used latrunculin B (Lat B) to depolymerize F-actin and jasplakinolide (Jasp) to stabilize and promote actin filament assembly. By comparison, TDG treatment produced a decrease in stress fiber density comparable to ROCK inhibition.

**Figure 5.**
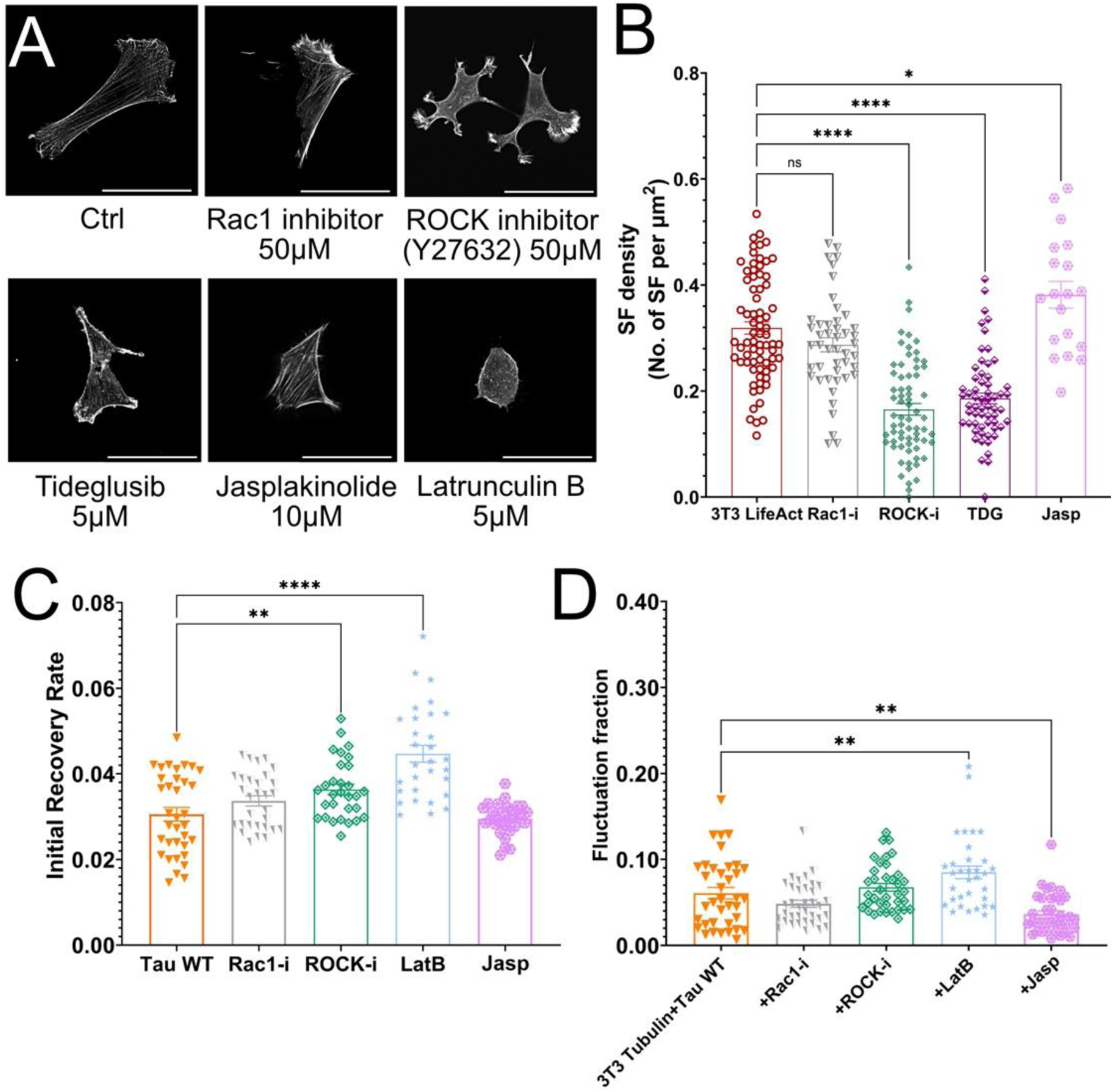
Actin cytoskeletal state influences Tau turnover without proportionally altering microtubule fluctuations. (A) Representative fluorescence images of 3T3 LifeAct cells following treatment with actin-regulatory drugs, illustrating changes in stress fiber organization and cell morphology. Scale bar: 50 µm. (B) Quantification of stress fiber density (number of stress fibers per μm²) in control and drug-treated cells. *3T3 LifeAct: n = 73; Rac1-i: p = 0.2222, n = 46; ROCK-i: p < 0.0001, n = 68; TDG: p < 0.0001, n = 60; Jasp: p = 0.0324, n = 20.* Data are presented as mean ± SEM (ns p > 0.05, *p < 0.05, **p < 0.01, ***p < 0.001, ****p < 0.0001). (C) Initial FRAP recovery rate of EGFP-Tau WT following treatment with Rac1-i, ROCK-i, latrunculin B, or jasplakinolide. *Tau WT: n = 35; Rac1-i: p = 0.1329, n = 30; ROCK-i: p = 0.0023, n = 30; Lat B: p < 0.0001, n = 30; Jasp: p > 0.9999, n = 30*. (D) Microtubule fluctuation fraction measured in 3T3 cells co-expressing tubulin and Tau WT under the same treatment conditions. *Tau WT: n = 37; Rac1-i: p = 0.3237, n = 36; ROCK-i: p = 0.8848, n = 37; Lat B: p = 0.0040, n = 33; Jasp: p = 0.0024, n = 36*.

Next, we quantified Tau turnover using FRAP in 3T3 cells expressing WT Tau. Disruption of actin produced consistent changes in Tau binding dynamics. Treatment with latrunculin B resulted in an increase in the initial FRAP recovery rate of Tau (Fig. 5C). Similarly, ROCK inhibition also increased Tau recovery rate compared to untreated cells. Notably, the change in Tau dynamics in ROCK-inhibited cells occurred without corresponding increases in microtubule network fluctuations (Fig. 5D), further supporting the idea that Tau exchange rates can be uncoupled from the dynamic state of the underlying tubulin network. In contrast, jasplakinolide-treated cells exhibited opposing effects on microtubule fluctuation fraction with actin stabilization decreasing fluctuation levels. These results suggest that Tau binding dynamics are sensitive to changes in F-actin organization, potentially reflecting a mechanosensitive component that integrates actin-dependent contractility and cell geometry with local microtubule structure.

### GSK-3β inhibition affects membrane poration and actin architecture

To test whether the protective effect of TDG against stretch-induced membrane poration arises from changes in actin organization and consequential changes in cell viscoelastic behaviour, Tau WT-expressing cells were subjected to AFM-based force-clamp measurements following treatment with actin-targeting agents (Fig. 6A-D) (see Supplementary Fig. 5 and Supplementary Text for results in 3T3 WT cells). In Tau WT cells, TDG treatment and ROCK inhibition produced modest reductions in cell stiffness relative to untreated Tau-expressing controls, whereas actin depolymerization with latrunculin B caused a pronounced decrease in stiffness, and actin stabilization with jasplakinolide had little effect.

**Figure 6.**
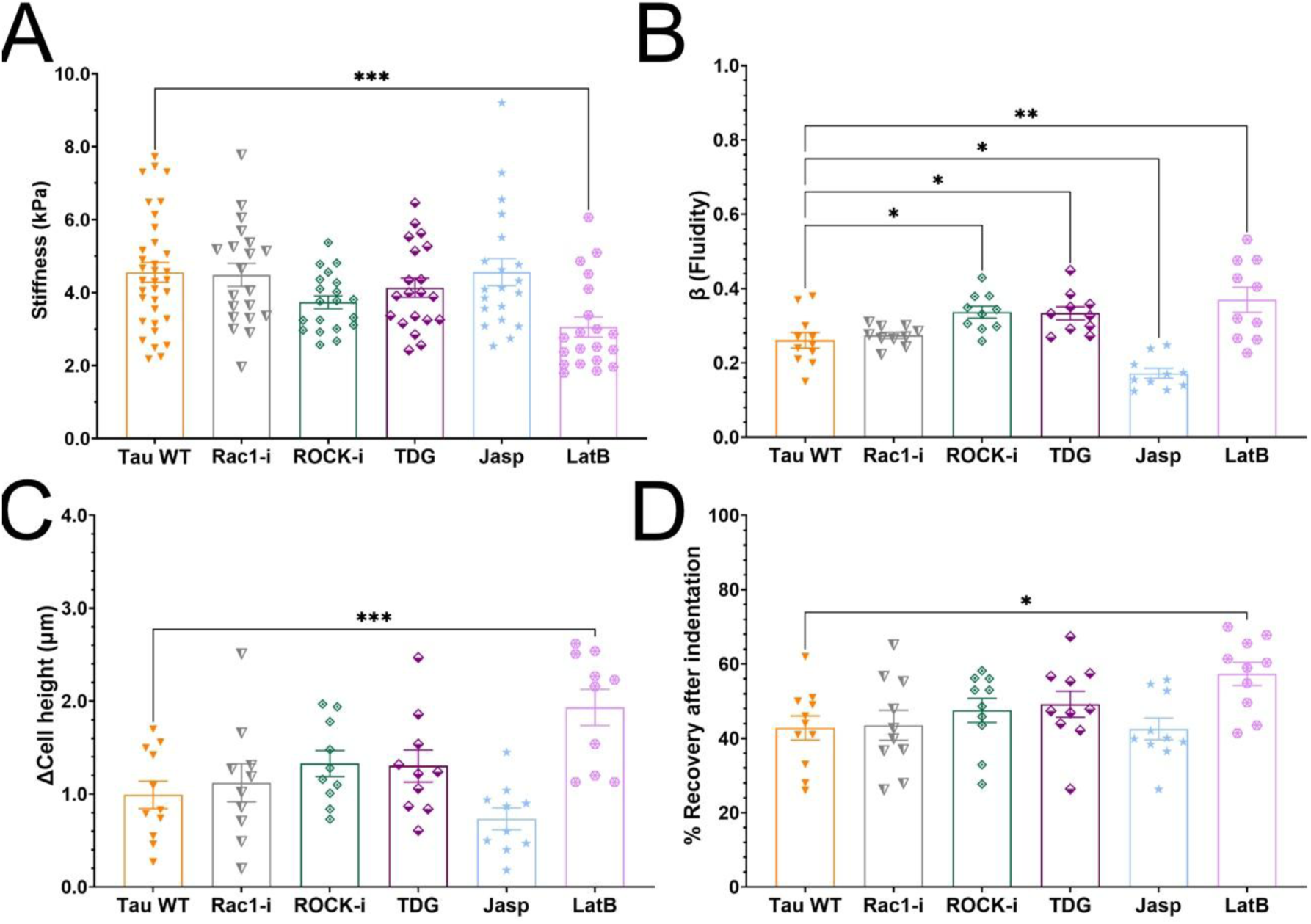
Pharmacological perturbation of actin organization alters cell morphology and mechanical properties in Tau WT 3T3 cells. (A) Effective cell stiffness of Tau WT 3T3 cells measured by AFM indentation following treatment with Rac1 inhibitor (Rac1-i), ROCK inhibitor (ROCK-i), TDG, jasplakinolide (Jasp), or latrunculin B (Lat B). *Tau WT: n = 34; Rac1-i: p > 0.9999, n = 20; ROCK-i: p = 0.1438, n = 20; TDG: p = 0.7487, n = 20; Jasp: p > 0.9999, n = 20; Lat B: p = 0.0008, n = 20.* Data are presented as mean ± SEM (ns p > 0.05, *p < 0.05, **p < 0.01, ***p < 0.001, ****p < 0.0001). (B) Power-law fluidity exponent β extracted from AFM force-relaxation measurements. *Tau WT: n = 11; Rac1-i: p = 0.9873, n = 10; ROCK-i: p = 0.0369, n = 10; TDG: p = 0.0490, n = 10; Jasp: p = 0.0107, n = 10; Lat B: p = 0.0012, n = 10*. (C) Creep deformation following pharmacological perturbation. *Tau WT: n = 11; Rac1-i: p = 0.9726, n = 10; ROCK-i: p = 0.4616, n = 10; TDG: p = 0.5382, n = 10; Jasp: p = 0.7052, n = 10; Lat B: p = 0.0007, n = 10*. (D) Percent recovery after indentation, reflecting viscoelastic recovery. *Tau WT: n = 11; Rac1-i: p > 0.9999, n = 10; ROCK-i: p = 0.7775, n = 10; TDG: p = 0.5296, n = 10; Jasp: p > 0.9999, n = 10; Lat B: p = 0.0130, n = 10*.

**Figure 7.**
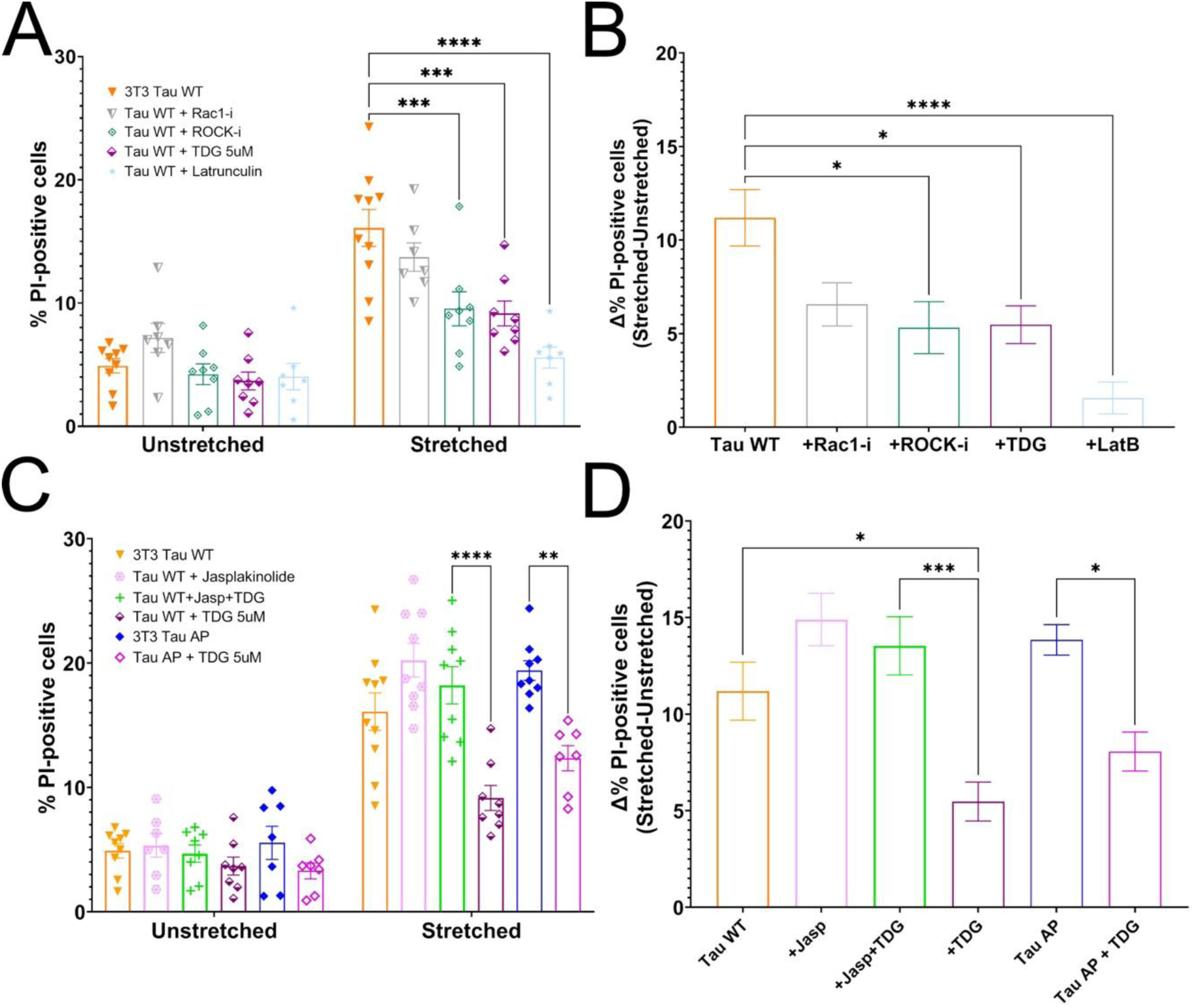
Actin cytoskeletal perturbations modulate Tau-dependent stretch-induced membrane poration. (A) Percentage of PI-positive cells in Tau WT-expressing 3T3 cells under unstretched and stretched conditions following treatment with Rac1 inhibitor (Rac1-i), ROCK inhibitor (ROCK-i), TDG, or latrunculin B (Lat B). *Unstretched 3T3 Tau WT: n = 9; Rac1-i: p = 0.8575, n = 7; ROCK-i: p = 0.9999, n = 8; TDG: p = 0.9936, n = 8; Lat B: p = 0.9994, n = 7 | Stretched 3T3 Tau WT: n = 10; Rac1-i: p = 0.8107, n = 7; ROCK-i: p = 0.0010, n = 8; TDG: p = 0.0004, n = 8; Lat B: p < 0.0001, n = 7.* Data are presented as mean ± SEM (ns p > 0.05, *p < 0.05, **p < 0.01, ***p < 0.001, ****p < 0.0001). (B) Normalized relative %PI uptake, calculated by subtracting the mean %PI of unstretched controls to reflect strain-induced changes for Tau WT-expressing cells across treatment conditions. *Rac1-i: p = 0.0910; ROCK-i: p = 0.0125; TDG: p = 0.0159; Lat B: p < 0.0001*. (C) PI positivity in unstretched and stretched Tau WT-expressing cells treated with jasplakinolide (Jasp), TDG, or combined jasplakinolide + TDG. *Unstretched 3T3 Tau WT: n = 9; Jasp: p > 0.9999, n = 9; Jasp + TDG: p > 0.9999, n = 9; TDG: p = 0.9997, n = 8; Tau AP: p > 0.9999, n = 7; Tau AP + TDG: p = 0.9969, n = 7 | Stretched 3T3 Tau WT: n = 10; Jasp: p = 0.1751, n = 9; Jasp + TDG: p = 0.9463, n = 9; TDG: p = 0.0007, n = 8; Tau AP: p = 0.4982 n = 9; Tau AP + TDG: p = 0.4109, n = 7. TDG vs. Jasp + TDG: p < 0.0001. Tau AP vs. Tau AP + TDG: p = 0.0015*. (D) Corresponding change in PI positivity highlighting differential effects of actin stabilization and Tau dephosphorylation. *Jasp: p = 0.2759; Jasp + TDG: p = 0.7473; TDG: p = 0.0266; Tau AP: p = 0.6365; Tau AP + TDG: p = 0.5400. TDG vs. Jasp + TDG: p = 0.0008. Tau AP vs. Tau AP + TDG: p = 0.0392*.

We next quantified time-dependent viscoelastic behaviour under sustained loading. TDG treatment and ROCK inhibition were associated with increased power-law fluidity exponent β and increased creep deformation, although the latter changes were not statistically significant. Rac1 inhibition did not meaningfully alter stiffness, β, creep, or recovery. Stabilization of filamentous actin with jasplakinolide decreased β, whereas actin depolymerization with latrunculin B increased β, creep deformation, and post-load recovery. These results suggest that disruption of Rho/ROCK-dependent actomyosin mechanics or complete actin depolymerization pushes cells toward a softer, more fluid-like state, indicating that TDG-associated protection is linked to an actin-mediated effect that is not fully counteracted by Tau-dependent microtubule reinforcement.

### Actin depolymerization protects against stretch-induced poration

To test the effect of actin organization on membrane vulnerability under rapid mechanical loading, we examined how pharmacological perturbations of the actin cytoskeleton affected stretch-induced membrane poration in Tau-expressing 3T3 cells. Cells were treated with Rac1 inhibitor, TDG, ROCK inhibitor, or latrunculin B prior to mechanical stretch. Under unstretched conditions, PI uptake remained low across all groups. Following high-rate stretch, ROCK inhibition and TDG treatment reduced stretch-induced PI uptake, with latrunculin B producing the strongest decrease. Rac1 inhibition had no significant effect, consistent with its limited impact on stress-fiber organization. We next examined whether actin stabilization alters susceptibility to mechanical injury. Tau WT-expressing cells were treated with jasplakinolide, either alone or in combination with TDG, and Tau AP-expressing cells were treated with TDG. PI uptake remained low and unchanged under unstretched conditions across all groups. Jasplakinolide treatment alone did not significantly affect PI uptake following stretch, indicating that additional actin stabilization does not exacerbate membrane poration. However, jasplakinolide blunted the protective effect of TDG, suggesting that TDG-mediated protection requires actin remodelling rather than acting solely through Tau-dependent pathways. Furthermore, TDG reduced stretch-induced PI uptake in Tau AP-expressing cells, demonstrating that actin-mediated mechanical changes can override the effects of stabilized microtubules.

## Discussion

Cells experience mechanical loading at a range of timescales, but rapid deformation presents a unique challenge because strain can outpace the viscoelastic relaxation needed to dissipate internal stress. Under these conditions, injury susceptibility depends not only on the magnitude of applied strain but also on the cytoskeleton’s ability to reorganize and redistribute load. Our results indicate that Tau overexpression fundamentally alters this balance by shifting the cytoskeleton toward a more rigid and less adaptive mechanical state, which in turn increases the likelihood of membrane poration during high strain-rate stretch (Fig. 8). Tau’s canonical role as a microtubule-associated protein that stabilizes microtubules is well established (24, 54), and its binding affinity is regulated by multisite phosphorylation (46, 55). Our results show that the expression of Tau in a non-neuronal cell type was sufficient to increase susceptibility to stretch-induced membrane poration. Tau WT and the dephospho-mimetic Tau AP variant produced the largest increases in cell stiffness and stretch-induced poration, as well as the strongest suppression of microtubule fluctuations; this was consistent with enhanced microtubule binding affinity and reduced filament mobility in these variants (41, 45, 56, 57). These findings suggest that while microtubule stabilization may support long-timescale processes such as intracellular transport and structural maintenance, excessive stabilization can increase vulnerability to acute mechanical insult by restricting the rapid cytoskeletal rearrangements required to dissipate load. It has previously been suggested that tau intermediate aggregates may promote neuronal damage through membrane disruption (58).

**Figure 8.**
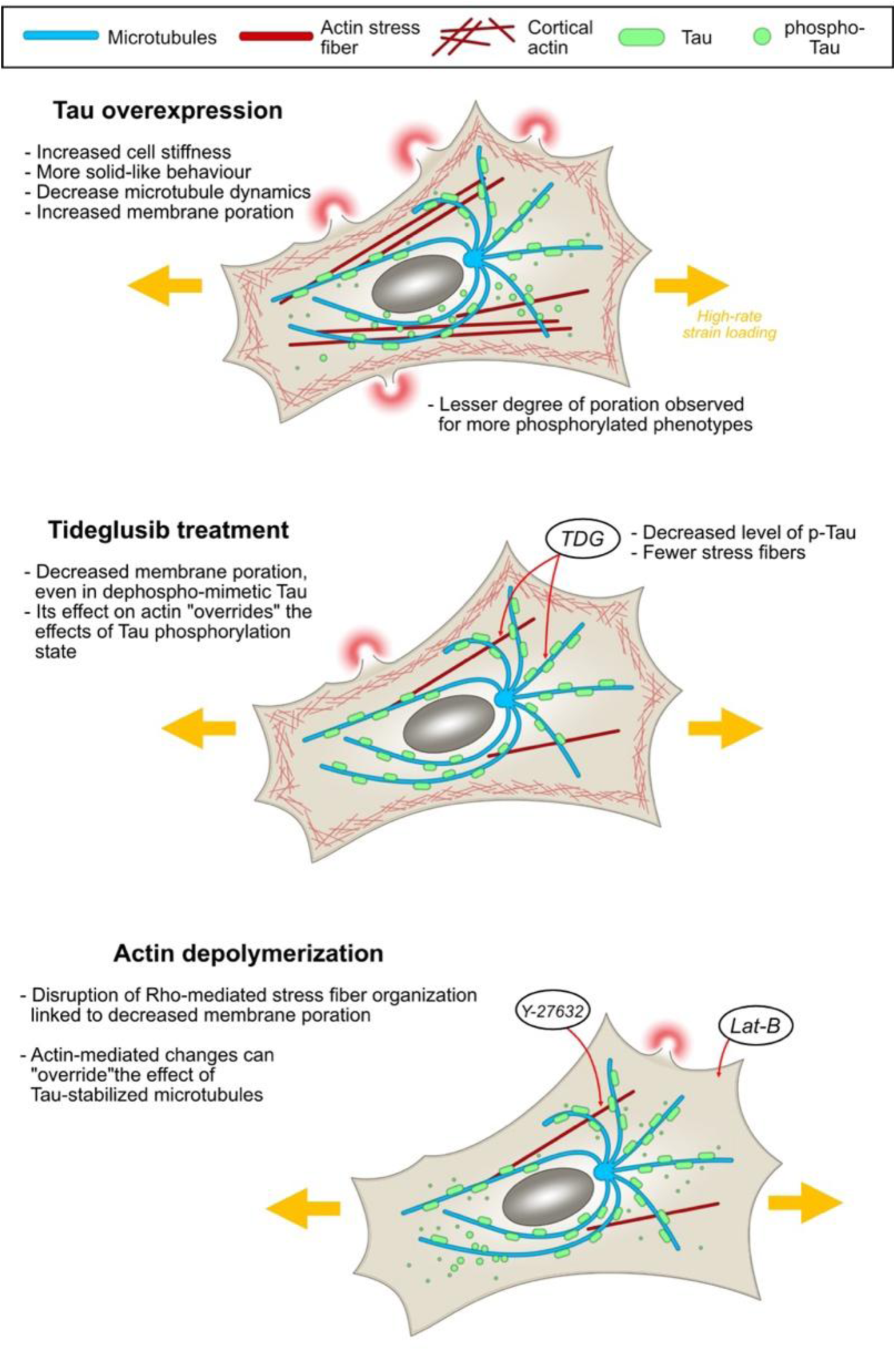
Graphical summary of the results presented in this study.

Within tensegrity-based models of cell mechanics (59, 60), an over-stabilized microtubule network behaves as an overly rigid internal scaffold. This rigidity would prevent the deformation load from being distributed normally and turn them into localized stress concentrations at the membrane-cortex interface, increasing the likelihood of membrane poration rather than adaptive viscoelastic deformation. Therefore, rather than serving as a universally protective stabilizer, increased microtubule stabilization shifts the cytoskeleton toward a less adaptable configuration. Previous experimental and modelling studies demonstrate that networks unable to rapidly reorganize exhibit stress stiffening and mechanical failure under high loading rates (61, 62). In Tau-expressing cells, this manifests as increased stiffness, reduced cytoskeletal fluidity, and heightened susceptibility to stretch-induced membrane poration. This is consistent with in vitro studies showing that Tau forms cooperative assemblies on compacted microtubule lattices and that lattice expansion or strain destabilizes these assemblies (63). These mechanically driven changes likely alter Tau-microtubule interactions during rapid deformation, feeding back into the cytoskeletal mechanical state captured in our experiments.

Because GSK-3β phosphorylates Tau, its inhibition was expected to decrease Tau phosphorylation and further stabilize microtubules. Indeed, TDG treatment reduced phospho-Tau levels and suppressed microtubule fluctuations. However, TDG unexpectedly reduced stretch-induced membrane poration as well. This apparent paradox may be related to TDG’s off-target effect on the organization of the actin cytoskeleton. The protective effect therefore appears to arise from actin reorganization rather than microtubule behaviour alone. These observations suggest that membrane vulnerability in this system is governed not by microtubule stability alone, but by the interaction between Tau-dependent microtubule stiffening and actin architecture, which together determine cytoskeletal-membrane force transmission. For example, actin stabilization with jasplakinolide did not affect poration on its own but eliminated the protective effects of TDG when applied concurrently, hinting that actin flexibility, more than filament abundance, is essential for load dissipation. Furthermore, our results indicate that Tau-microtubule interactions are sensitive to changes in the F-actin network; disrupting stress fibers with latrunculin B or ROCK inhibition reduced stiffness and decreased membrane poration in Tau-expressing cells. This suggests that Tau dynamics may be influenced by actin-dependent contractility and cell geometry. In this view, Tau integrates mechanical inputs from multiple cytoskeletal components, linking mechanical conditions in the actin network to local microtubule structure. This interpretation aligns with emerging evidence that Tau participates in broader mechanotransduction pathways, including those regulating cytoskeletal tension and nuclear mechanics (64, 65). In this context, we speculate that Tau may function as a mechanically responsive element that respond to both microtubule state and actin networks to shape how cells distribute and tolerate rapid deformation.

Our findings align with prior work demonstrating that membrane vulnerability to mechanical loading is regulated by cytoskeletal organization and viscoelastic relaxation dynamics. Prado and LaPlaca (66) showed that traumatic loading induces transient membrane disruptions in neurons and that the extent of poration is strongly dependent on loading rate, reflecting the limited time available for cytoskeletal and membrane relaxation under rapid strain. They further reported that manipulating the actin cortex modulates both permeability and resealing, which parallel our finding that disruption of actin stress fibers in Tau-expressing fibroblasts reduces stretch-induced PI uptake. Although our experiments used a non-neuronal system, the strain-rate-dependent injury phenotype resembles observations in neuronal stretch-injury models (67, 68). These findings may have relevance for mild traumatic brain injury (mTBI), where axons, whose functions depend heavily on dynamic microtubules, experience rapid deformation. Stabilized microtubules may limit the rapid cytoskeletal remodelling necessary to protect the membrane, potentially amplifying initial mechanical damage. Conditions that hyper-stabilize microtubules, whether via Tau overexpression or altered phosphorylation may increase vulnerability to acute mechanical loading by suppressing cytoskeletal fluidity. It must also be noted that both Tau dysregulation and actin pathology characterize neurodegeneration. Tauopathies involve loss of normal Tau-microtubule interactions (69) followed by compensatory hyper-stabilization of remaining microtubules (70, 71), producing networks that are structurally rigid but poorly adaptable. Actin pathology, including formation of cofilin-actin rods observed in neurodegenerative diseases (72), similarly reduces cytoskeletal flexibility. The convergence of microtubule and actin rigidity may therefore represent a mechanical vulnerability shared across acute injury and chronic neurodegeneration. In another clinical context, these findings may have implications for conditions such as chemotherapy-induced peripheral neuropathy. Microtubule-stabilizing chemotherapeutics such as paclitaxel may exacerbate mechanical vulnerability by shifting the cytoskeleton toward the rigid, low-fluidity state described here. Our results also highlight that pharmacological agents such as tideglusib, intended to modulate Tau phosphorylation, can exert off-target effects on other cytoskeletal systems including the actin network, potentially producing unintended mechanical consequences. Targeting the interplay between Tau-microtubule interactions and actin flexibility may therefore represent a promising therapeutic approach for reducing acute mechanical injury while avoiding long-term cytoskeletal dysfunction.

## Materials and Methods

### Cell culture

NIH 3T3 mouse fibroblasts were cultured in Dulbecco’s modified Eagle’s medium (DMEM, Gibco) supplemented with 10% heat-inactivated fetal bovine serum (Gibco), 100 U/mL penicillin, 100 µg/mL streptomycin (Gibco), and 2 mM L-glutamine. Cells were maintained at 37 °C in a humidified Thermo Scientific 3110 CO₂ incubator under a 5% CO₂ atmosphere.

### Cloning and generation of stable cell lines

mCherry-Tubulin-6 was a gift from Michael Davidson (Addgene plasmid #55147; http://n2t.net/addgene:55147; RRID:Addgene_55147). pRK5-EGFP-Tau constructs were gifts from Karen Ashe, including wild-type Tau (Addgene plasmid #46904; http://n2t.net/addgene:46904; RRID:Addgene_46904), Tau P301L (Addgene plasmid #46908; http://n2t.net/addgene:46908; RRID:Addgene_46908), Tau E14 (Addgene plasmid #46907; http://n2t.net/addgene:46907; RRID:Addgene_46907), and Tau AP (Addgene plasmid #46905; http://n2t.net/addgene:46905; RRID:Addgene_46905). DNA encoding EGFP-Tau, EGFP-Tau E14, EGFP-Tau AP, or EGFP-Tau P301L was amplified by PCR and inserted into the lentiviral vector pHR using Gibson assembly. Lentivirus production was performed by co-transfecting the pHR constructs with the helper plasmids P8.91 and PMD2.G into HEK293T cells using Transit 293 (Mirus). Viral supernatants were collected 48 h post-transfection, passed through a 0.4 µm filter, and used directly to infect wild-type NIH 3T3 cells. Expression of the EGFP-tagged constructs in infected cells was verified by fluorescence microscopy, and only EGFP-positive cells were selected for subsequent analysis.

### Cell Stretching System

High-strain uniaxial stretching of cells was performed using a custom in-house system based on a voice coil actuator (Zaber X-DMQ12L-AE55D12), capable of precise cyclic or instantaneous loading within a cell culture incubator. Actuator control was implemented via Zaber Launcher software combined with custom MATLAB scripts. Detailed schematics and operation procedures are described in (73). PDMS wells for cell culture were prepared from Sylgard™ 184 slabs with integrated square wells (10 mm × 10 mm) and bottom sealing layers. Wells were sterilized by UV, coated with polydopamine (15 wt/vol%), and washed with PBS prior to use. Full fabrication and coating procedures, including dimensions, curing steps, and coating conditions, are described in (73).

### Microscopy

Fluorescence images were acquired using a Nikon Ti2 inverted fluorescence microscope equipped with a Crest V3 spinning disk confocal system. Plan Apo λD 20x objective or Plan Apo VC 60× water-immersion DIC N2 objective was used depending on the experiment, and image capture was performed with a Kinetix camera.

### Membrane Damage Assay

Membrane integrity was assessed using Hoechst (Abcam, Cambridge, UK, ab228551) and propidium iodide (PI, BioShop Canada, Burlington, ON, Canada, PPI888.5) staining following 1-hour exposure to test reagents and mechanical stretching. After stretching, cells were incubated an additional 30 min, then imaged on the previously described microscope setup (See: Microscopy) with the 20x objective. PI-positive cells were quantified as a percentage of total nuclei. In GFP-Tau expressing lines, non-expressing cells were excluded based on GFP signal. Detailed protocols for staining, imaging, and quantitative analysis are provided in (73).

### Pharmacological treatments

Cells were plated at a density of 30,000 cells/cm² and cultured for 3 days in differentiation medium, after which they were exposed to the test compounds for 1 h. For pharmacological treatments, the following drug concentrations were used: 1 µM paclitaxel (BioShop Canada, PAX960), 5 µM tideglusib (Sigma, SML0339), 10 µM jasplakinolide (Abcam, 141409), 50 µM ROCK inhibitor Y-27632 (Abcam, 120129), 50 µM Rac1 inhibitor (Sigma, Z62954982), and 5 µM latrunculin B (Abcam, 144291). Stock solutions of each compound were prepared in dimethyl sulfoxide (DMSO; Invitrogen, D12345) and diluted to the final concentration in DMEM immediately prior to use.

### AFM measurements

Creep experiments were performed using an in-house built atomic force microscope (AFM) (40) equipped with a pyramidal-tipped cantilever (PNP-TR-Au Cant. 2, NanoWorld). Prior to each measurement, the cantilever deflection sensitivity was calibrated using a hard-substrate indentation procedure. The nominal spring constant provide by the manufacturer was used (0.08 N/m) (74).

The cantilever was then positioned above an isolated cell and lowered at a constant approach velocity (3.5 µm/s) until a pre-defined set point force (*F*_0_) of 2nN was reached.

### AFM data analysis

To ensure robust contact-point detection and model fitting, force-distance curves were screened for data quality prior to analysis. Curves displaying excessive noise, abrupt force fluctuations, or irregular features inconsistent with elastic indentation indicative of poor tip-sample contact or instrumental artefacts were discarded. The contact point was determined using a MATLAB script based on the algorithm developed by Lin et al. (2006), using least squares minimisation. For every point in the curve, a linear regression model and a Hertz model was fitted to the left and to the right of the point of interest respectively. The sum of the error of the two fits was calculated for every point, and the point with the least summed error was chosen as the contact point. The section of the curve to the right of the contact point was subsequently taken to be the post-contact region. The initial cell height was determined from the cantilever’s vertical position at the point of first contact. Briefly, for each cell measurement, a corresponding measurement was collected on the glass surface in a region near the cell perimeter. Given that the cell and glass measurements have the same starting point of piezo-displacement z = 0, the cell height h may be calculated as the difference between the z values of the contact points (*z*_*cp*_) from the respective curves.

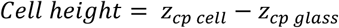

To determine the cell elastic modulus from the force-distance curve, the post-contact region of the f-d curve was fitted with the Bilodeau model for pyramidal contact to determine the Young’s modulus of the cell, E, using the following equation (75, 76):

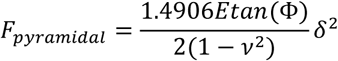

(F=force, δ=indentation depth, ν=Poisson’s ratio, **Φ**=opening angle of the pyramidal tip)

To ensure the analysis is performed on the pertinent section of the non-linear region, the fitting was limited to segment of the curve where F<1.5nN

After establishing contact, the cantilever was loaded to a pre-set constant force (*F*_0_) of 2nN, at which point a 60-s force-clamp (constant-force hold) was initiated. During this period the AFM continuously recorded the cantilever height with a sampling interval of 0.01 s. The gradual increase in indentation (δ(*t*)) during the hold reflects time-dependent creep deformation of the cell. The creep compliance *J*(*t*) was computed from the indentation history using the pyramidal-contact relation:

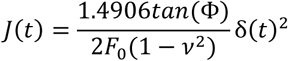

The instantaneous indentation was computed as the difference between the cantilever Z-piezo height at each time point and the initial cell height. At the end of the 60-s creep period, the final cantilever height was recorded, yielding the cell height after creep. The cantilever was then fully retracted to remove the load, and the cell was allowed to recover for 10 s. After this recovery period, a second approach was performed to determine the cell height after recovery.

### Quantification of Viscoelastic Parameters

The experimentally derived creep compliance (*J*(*t*)) was fitted with a power-law rheological model,

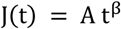

where β is the fluidity parameter, characterizing the degree to which the cell behaves as a fluid-like versus solid-like material. The fitting was performed across the full 60 s creep interval for each cell. Additionally, the indentation depth was defined as

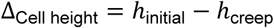

where ℎ_initial_ is the initial cell height at first contact and ℎ_creep_ is the height after 60 s of creep under constant load. After the 10-s recovery interval, the percent recovery was computed as

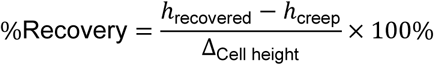

where ℎ_recovered_ is the cell height measured at re-contact following recovery.

### Live-cell microtubule fluctuation imaging

Fluorescence time-lapse imaging was performed on the previously described microscope setup (See: Microscopy) with the 60x objective. Images were acquired every 2 s for a total duration of 120 s. Temporal dynamics of fluorescently labelled tubulin filaments were quantified using an image-based analysis method (77). For each image series, a custom ImageJ macro was used to calculate the standard deviation of fluorescence intensity at each pixel over time. The standard deviation was divided by the mean intensity to generate a coefficient of variation (CoV) map, which was then normalized to the maximum intensity in the dataset. Bleaching correction was applied using the Bleach Correction plugin in ImageJ (78) prior to analysis to minimize contributions from non-dynamic signal changes. To focus on filament dynamics, cytoplasmic regions were excluded by applying Li’s thresholding algorithm (79), and non-filament pixels were set to NaN. High-fluctuation areas were identified using an arbitrary threshold value, and the fraction of pixels above the threshold relative to the total number of pixels per cell was calculated within defined regions of interest (ROIs).

### Actin Stress Fiber Segmentation

NIH 3T3 cells expressing LifeAct-GFP were imaged on the previously described microscope setup (See: Microscopy) with the 60x objective to visualize the actin cytoskeleton. Acquired images were segmented using the FAST (Filament Analysis and Segmentation Tool) (53) application which identifies cytoskeletal structures based on filamentous morphology. Thread-like actin structures were considered as actin stress fibers. Segmentation results were visually inspected to confirm accurate identification of stress fibers (see Supplementary Fig. 4). Stress fiber counts were normalized to cell area to obtain the density of stress fibers per unit area.

### Fluorescence Recovery After Photobleaching (FRAP)

Cells expressing EGFP-tagged Tau constructs were cultured on glass-bottom dishes and imaged at 37 °C in phenol red-free imaging medium. Prior to FRAP experiments, cells were allowed to equilibrate on the microscope stage for at least 10 min to minimize thermal or mechanical drift. Only well-spread, isolated cells with clear filamentous Tau localization along microtubules were selected for analysis. FRAP experiments were performed on the previously described microscope setup (See: Microscopy) with the 60x objective. A small circular region of interest (ROI) was positioned along Tau-expressing microtubule bundle. Pre-bleach images were acquired for 10 frames at low laser power to establish baseline fluorescence. Photobleaching was achieved by briefly exposing the ROI to high laser power (100% power for 1s), resulting in an instantaneous reduction of fluorescence intensity within the bleached region. Following bleaching, fluorescence recovery was monitored by time-lapse imaging at low laser power. Images were acquired at rate of 1 fps until the fluorescence approached a plateau. Fluorescence intensity within the bleached ROI was quantified at each time point and corrected for background signal measured in a cell-free region. To account for overall photobleaching during image acquisition, intensities were normalized to a non-bleached reference region within the same cell. Recovery curves were further normalized such that the mean pre-bleach intensity was set to 1 and the first post-bleach intensity to 0. To quantify Tau turnover dynamics, a recovery rate was calculated by performing a linear regression on the initial portion of the normalized FRAP recovery curve; fluorescence intensity values obtained during the first 5 s following photobleaching were fitted with a straight line, and the resulting slope was defined as the recovery rate. This metric captures the early-phase recovery kinetics and provides a model-independent measure of Tau exchange at microtubules.

### Quantification of Phospho-Tau Immunofluorescence Intensity

Phospho-Tau signal intensity was quantified from immunofluorescence images to assess changes in Tau phosphorylation following pharmacological treatment. The cells were fixed and stained with phospho-Tau antibody (primary Ab: Invitrogen, MN1020; secondary Ab: goat anti-rabbit IgG H&L [Alexa Fluor (Thermo Fisher Scientific, Waltham, MA, USA) 647], Abcam, ab150079) and imaged on the previously described microscope setup (See: Microscopy) with the 60x objective. Only cells expressing EGFP-tagged Tau were included in the analysis. We found that chemical fixation can cause partial dissociation of Tau from microtubules; therefore the fluorescence measurements primarily reflect the total cellular phospho-Tau signal rather than microtubule-bound Tau specifically.

Image analysis was performed using Fiji/ImageJ (NIH). For each image, individual cells were manually outlined based on the EGFP-Tau channel to generate regions of interest (ROIs) corresponding to Tau-expressing cells. Mean phospho-Tau fluorescence intensity within each ROI was measured from the phosphor-Tau antibody channel. Background fluorescence was estimated from a cell-free region within the same field of view and subtracted from all measurements. To account for variability in Tau expression levels between cells, these values were normalized to the corresponding mean EGFP-Tau intensity measured within the same ROI. Normalized values were pooled across multiple fields of view and independent experiments for each condition.

### Statistical analysis

GraphPad Prism 10 (GraphPad Software Inc., United States) was used for graph generation and statistical analyses. Data are presented as mean ± standard error of the mean (SEM). One-way ANOVA was used, followed by Tukey’s multiple comparisons post-hoc tests. Differences were considered statistically significant when p ≤ 0.05 (ns p > 0.05, *p < 0.05, **p < 0.01, ***p < 0.001, ****p < 0.0001).

## Acknowledgments

This research was enabled in part by support provided by Research Computing Services (https://carleton.ca/rcs) at Carleton University.

## Author Contributions

GK, ARH and OEP conceived and designed the study. GK performed the experiments, data analysis, and statistical evaluation. VA contributed to data analysis. GK, ARH and OEP wrote the manuscript. All authors contributed to data interpretation and manuscript revisions.

## Supplementary figures

**Supplementary Figure 1.**
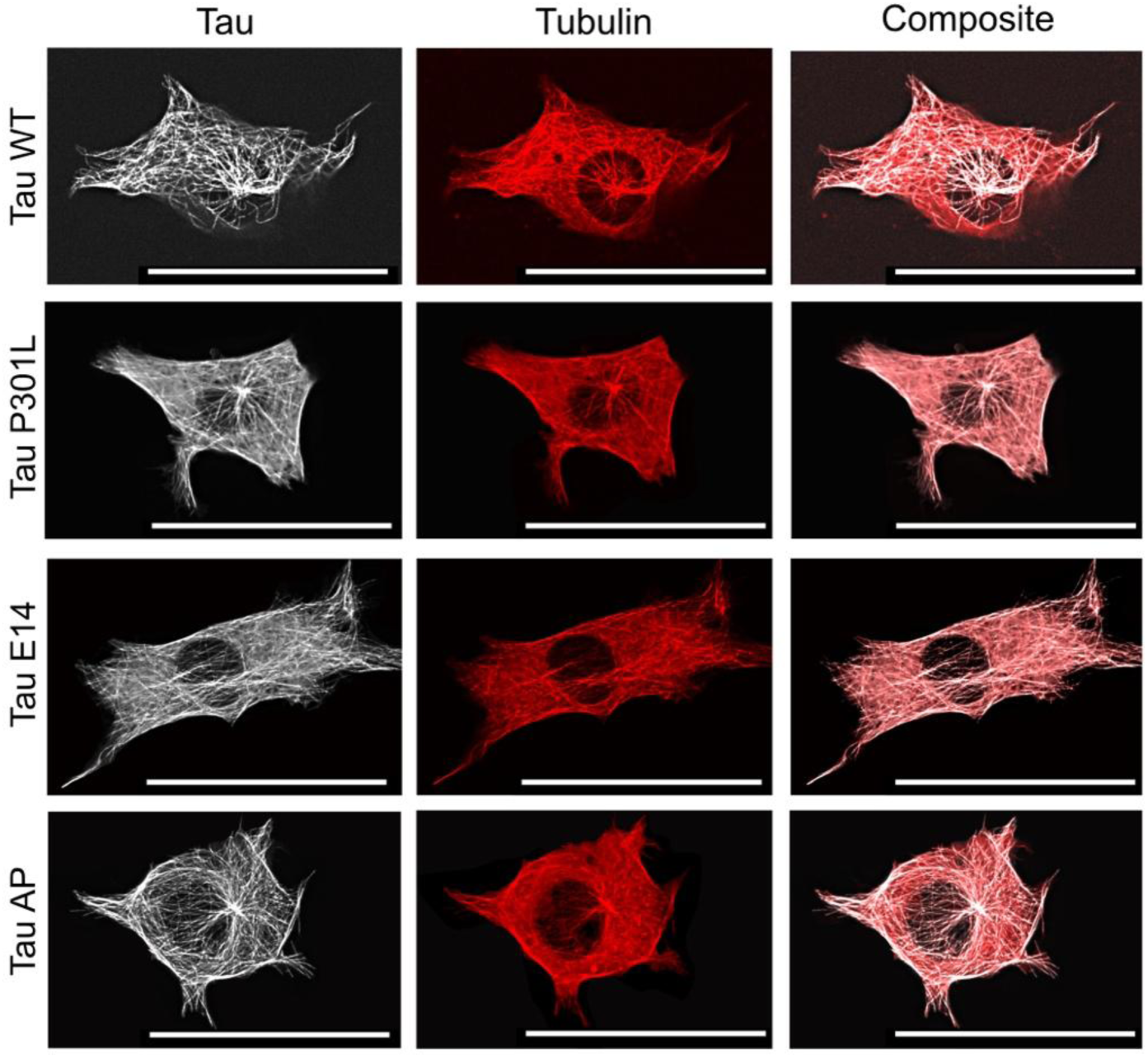
Tau localization relative to the microtubule network across Tau variants. Representative fluorescence images of 3T3 cells expressing EGFP-tagged Tau variants (WT, P301L, E14, and AP). Tau signal (left), tubulin (middle), and merged composite images (right) are shown for each condition. Across all variants, Tau exhibits a filamentous distribution that overlaps extensively with the microtubule network, with subtle differences in network organization and Tau enrichment apparent between constructs. Scale bar: 50µm.

**Supplementary Figure 2.**
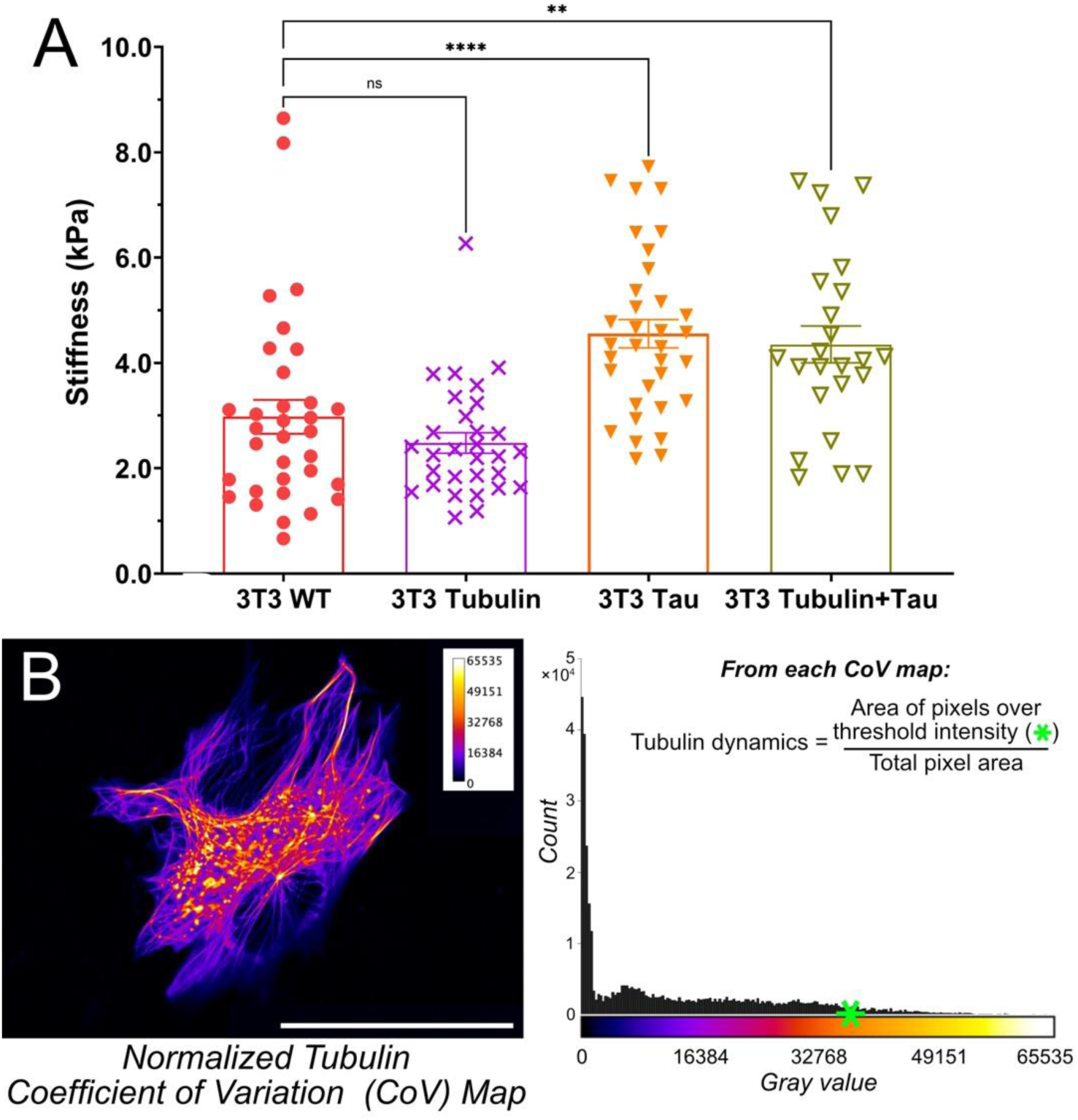
Effects of Tau expression on whole-cell stiffness and microtubule dynamics. (A) Cell stiffness measured by AFM indentation for 3T3 WT cells, tubulin-labeled controls, Tau-expressing cells, and cells co-expressing tubulin and Tau. Individual data points represent single-cell measurements; bars indicate mean ± SEM. Statistical significance is indicated as shown (ns, not significant; **p < 0.01; ****p < 0.0001; (3T3 Tubulin: p = 0.5734 vs. 3T3 WT). (B) Left: Representative example of a normalized tubulin coefficient of variation map. Scale bar: 50µm. Right: histogram of grayscale pixel intensity values from the thresholded cell area, with the tubulin dynamic calculated from the distribution as a metric of microtubule intensity fluctuations.

**Supplementary Figure 3.**
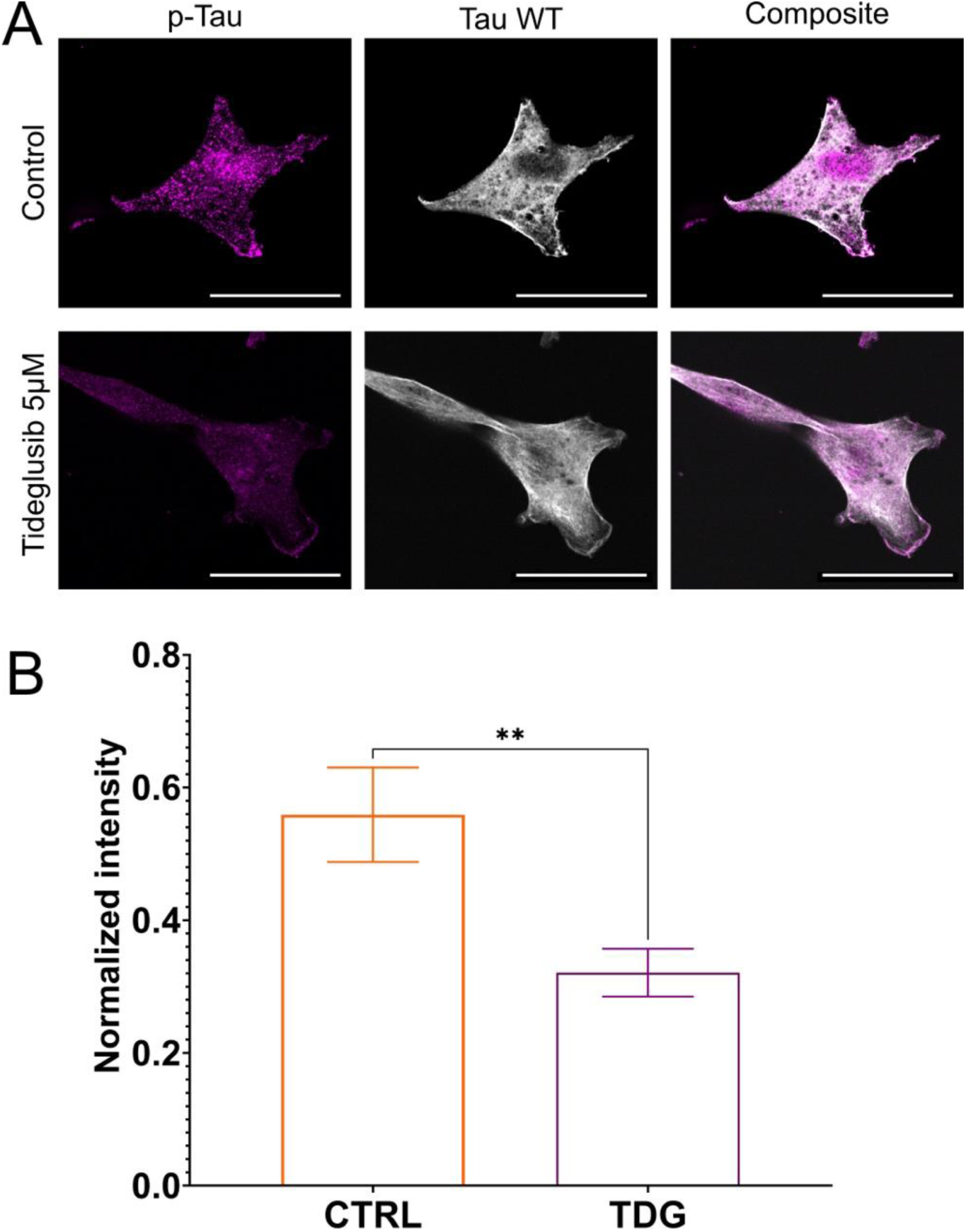
Tideglusib treatment reduces Tau phosphorylation without altering gross Tau distribution. (A) Representative fluorescence images of Tau WT-expressing 3T3 cells immunostained for phosphorylated Tau (p-Tau; magenta) (Stained with Invitrogen MN1020) and total Tau (grey) under control conditions and following treatment with 5 µM tideglusib. Composite images show reduced p-Tau signal intensity in tideglusib-treated cells. We found that chemical fixation can cause partial dissociation of Tau from microtubules; therefore the fluorescence measurements primarily reflect the total cellular phospho-Tau signal rather than microtubule-bound Tau specifically. Scale bar: 25µm. (B) Quantification of mean p-Tau fluorescence intensity per cell normalised to the mean Tau intensity demonstrates a significant reduction in p-Tau levels following tideglusib treatment compared to control. Bars represent mean ± SEM; **p < 0.01.

**Supplementary Figure 4.**
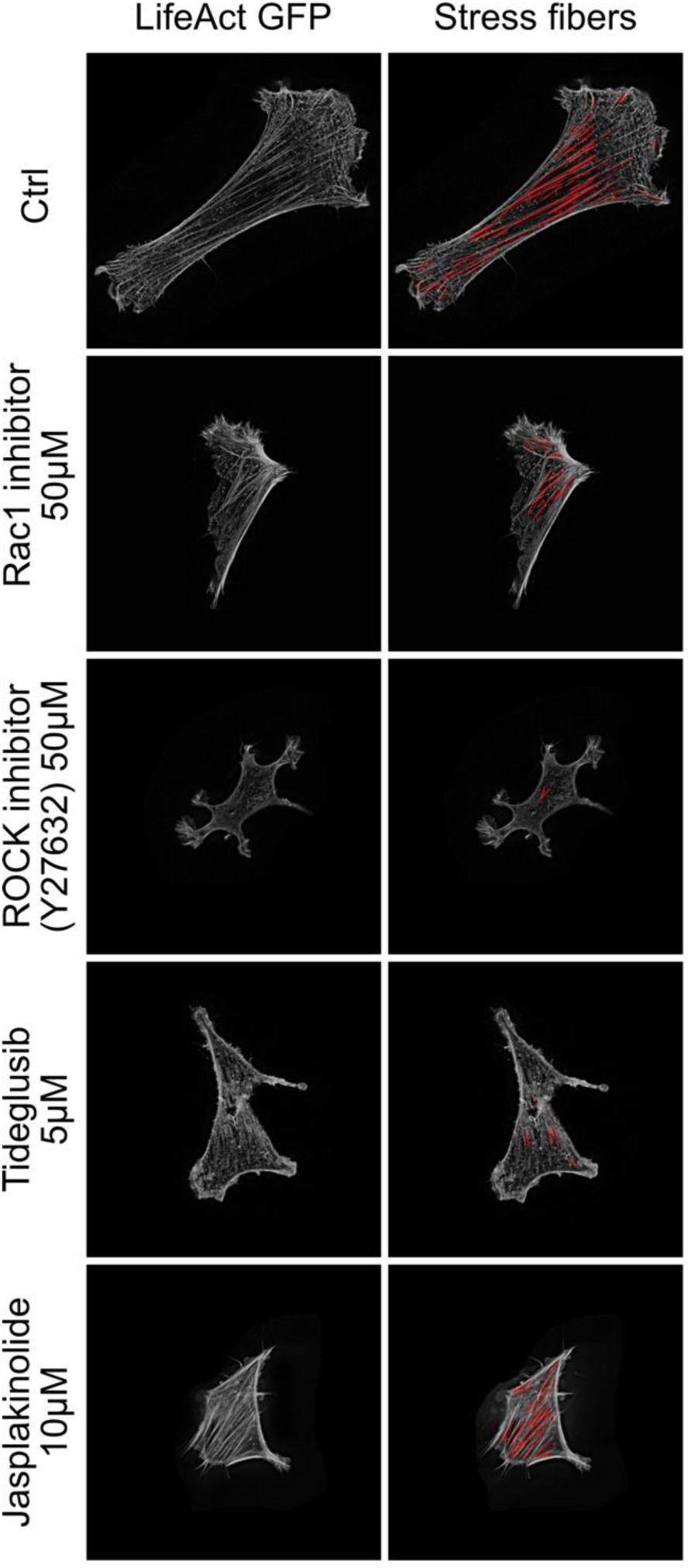
Representative examples demonstrating the FAST procedure used for stress fiber analysis.

**Supplementary Figure 5.**
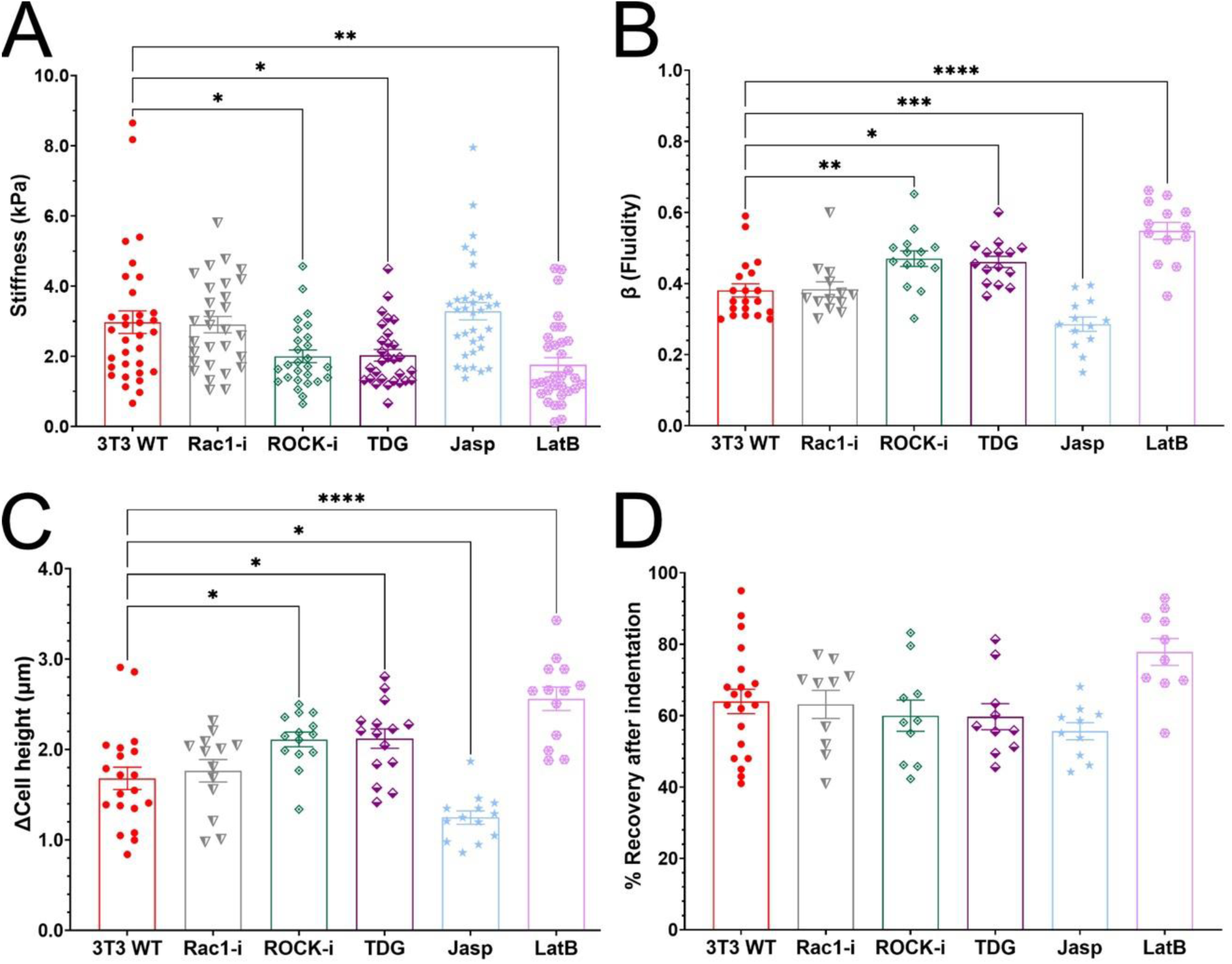
Pharmacological perturbation of actin organization alters cell morphology and mechanical properties in 3T3 WT cells. **(A)** Effective cell stiffness of 3T3 WT cells measured by AFM indentation following treatment with Rac1 inhibitor (Rac1-i), ROCK inhibitor (ROCK-i), TDG, jasplakinolide (Jasp), or latrunculin B (LatB). **(B)** Fluidity parameter (β) extracted from AFM force-relaxation measurements across treatment conditions. **(C)** Change in cell height following pharmacological perturbation. **(D)** Percent recovery after indentation, reflecting viscoelastic recovery behaviour. Data are shown as individual measurements with mean ± SEM; ns p > 0.05, *p < 0.05, **p < 0.01, ***p < 0.001, ****p < 0.0001.

## Supplementary text

To determine how pharmacological perturbations of actin organization modulate viscoelastic behaviour under sustained loading, cells were subjected to AFM-based force-clamp measurements following treatment with actin-targeting agents (Supplementary Fig. 4B-E). From these measurements, we quantified changes in the power-law fluidity exponent β, total creep deformation during the clamp, and post-loading recovery. Perturbations that reduced actomyosin contractility, including TDG treatment and ROCK inhibition, resulted in a significant increase in β relative to untreated controls (TDG: p = 0.0129; ROCK-i: p = 0.0087), indicating a shift toward more fluid-like behaviour. These treatments were also associated with increased deformation during the force clamp (TDG: p = 0.0281; ROCK-i: p = 0.0389). Despite these changes, recovery following load release was slightly reduced compared to control cells, although this effect did not reach statistical significance (p ≥ 0.9808 in both). In contrast, Rac1 inhibition did not significantly alter β, deformation, or recovery relative to untreated controls (p ≥ 0.9989 across parameters). Stabilization of filamentous actin with jasplakinolide produced a significant decrease in β (p = 0.0002) and creep deformation (p = 0.0448), consistent with a more solid-like mechanical response, while recovery was slightly reduced (p = 0.05521). Conversely, actin depolymerization with latrunculin B led to pronounced increases in β and deformation (both p < 0.0001), accompanied by a non-significant increase in recovery. These results imply that although multiple pathways contribute to actin regulation, Rho/ROCK-mediated stress fiber organization produces the most distinct shifts in cellular behaviour and may play a dominant role in determining membrane vulnerability under rapid mechanical loading. TDG appears to confer protection not through Tau phosphorylation but through a secondary action that disrupts Rho-dependent actin structures, reduces cell stiffness, and thereby limits stretch-induced membrane poration.

## Supplementary Video

Live-cell timelapse of fluorescent Tau labelled microtubules in untreated 3T3 Tau WT, paclitaxel (PTX) treated and tideglusib (TDG) treated cells. PTX and TDG treatments visibly reduce cytoskeletal fluctuations compared with control cells, highlighting the impact of microtubule stabilization and turnover inhibition on cellular dynamics.

